# Introgression of the *Triticum timopheevii* genome into wheat detected by chromosome-specific KASP markers

**DOI:** 10.1101/2022.05.04.490655

**Authors:** Julie King, Surbhi Grewal, Manel Othmeni, Benedict Coombes, Cai-yun Yang, Nicola Walter, Stephen Ashling, Duncan Scholefield, Jack Walker, Stella Hubbart-Edwards, Anthony Hall, Ian King

## Abstract

*Triticum timopheevii* (2n=28, A^t^A^t^GG) is a tetraploid wild relative species with great potential to increase the genetic diversity of hexaploid wheat *Triticum aestivum* (2n=42, AABBDD) for various important agronomic traits. A breeding scheme that propagated advanced backcrossed populations of wheat-*T. timopheevii* introgression lines through further backcrossing and self-fertilisation resulted in the generation of 99 introgression lines (ILs) that carried 309 homozygous segments from the A^t^ and G subgenomes of *T. timopheevii*. These introgressions contained 89 and 74 unique segments from the A^t^ and G subgenomes, respectively. These overlapping segments covered 98.9% of the *T. timopheevii* genome that has now been introgressed into bread wheat cv. Paragon including the entirety of all *T. timopheevii* chromosomes via varying sized segments except for chromosomes 3A^t^, 4G and 6G. Homozygous ILs contained between one and eight of these introgressions with an average of three per introgression line. These homozygous introgressions were detected through the development of a set of 480 chromosome-specific Kompetitive allele specific PCR (KASP) markers that are well-distributed across the wheat genome. Of these, 149 were developed in this study based on single nucleotide polymorphisms (SNPs) discovered through whole genome sequencing of *T. timopheevii*. A majority of these KASP markers were also found to be *T. timopheevii* subgenome specific with 182 detecting A^t^ subgenome and 275 detecting G subgenome segments. These markers showed that 98% of the A^t^ segments had recombined with the A genome of wheat and 74% of the G genome segments had recombined with the B genome of wheat with the rest recombining with the D genome of wheat. These results were validated through multi-colour in situ hybridisation analysis. Together these homozygous wheat-*T. timopheevii* ILs and chromosome-specific KASP markers provide an invaluable resource to wheat breeders for trait discovery to combat biotic and abiotic stress factors affecting wheat production due to climate change.

## 1 Introduction

The wild relatives of wheat provide genetic variation for most, if not all, traits of agronomic importance including resistances to abiotic stresses, e.g. *Thinopyrum bessarabicum* (King et al., 1997), biotic stresses, e.g. *Aegilops speltoides* (Niranjana et al., 2017) and *Aegilops ventricosa* (Cruz et al., 2016), novel variation for protein quality, e.g. *Thinopyrum intermedium* (Li et al., 2013) and mineral content, e.g. *Aegilops* sp. (Sharma et al., 2018) and increases in physiological traits such as photosynthetic capacity, e.g. *Thinopyrum elongatum* (Reynolds et al., 2001).

A wild relative that has proved to be of considerable interest due to the high levels of genetic variation it has been shown to contain is *Triticum timopheevii* subsp. *timopheevii* (Zhuk.) (2n=4x=28; A^t^A^t^GG). Traits of interest include resistance to leaf rust including the genes *Lr10, Lr18* and *Lr50* (Brown-Guedira et al., 2003; Leonova et al., 2010; McIntosh 1983; Singh et al., 2017), stem rust including the genes *Sr36, Sr37* and *Sr40* (Allard and Shands 1954; Dyck 1992; McIntosh and Gyarfas 1971), powdery mildew including the genes *Pm2, Pm6, Pm27* and *Pm37* (Allard and Shands 1954; Järve et al., 2000; Jørgensen and Jensen 1972; Perugini et al., 2008; Peusha et al., 1995; Qin et al., 2011), Fusarium head blight (Malihipour et al., 2016; Malihipour et al., 2017), black-point (Lehmensiek et al., 2004), Hessian fly, Septoria blotch, wheat curl mite and tan spot (Brown-Guedira et al., 1996). *T. timopheevii* has also been shown to be a source of resistance to abiotic stresses such as salt tolerance (Badridze et al., 2009; Yudina et al., 2016) and traits affecting grain quality such as milling yield and grain protein (Lehmensiek et al., 2006) and mineral content (Hu et al., 2017).

*Triticum timopheevii* belongs to the secondary gene pool of wheat as both genomes are very closely related to wheat although they show modifications. The A-genomes of durum wheat (*Triticum durum* Desf., 2n=4x=28; AABB), bread wheat (*Triticum aestivum* L., 2n=6x=42; AABBDD) and *T. timopheevii* are derived from *Triticum urartu* Thum. ex Gandil. (Dvořák et al., 1993) while the B-genomes of durum and bread wheat and the G-genome of *T. timopheevii* are derived from an *Ae. speltoides*-like species (Dvorak and Zhang 1990). A recent study showed that *Ae. speltoides* was not the direct donor of the G subgenome and diverged from the progenitor of the latter ca. 2.93 mya, although it was more closely related to it than to the B genome of wheat (Li et al., 2022). As in *T. durum* and *T. aestivum, T. timopheevii* has been shown to carry the 4A^t^L/5A^t^L translocation, but in addition, also contains a 6A^t^/1G/4G cyclic translocation (Jiang and Gill 1994; Maestra and Naranjo 1999; Salina et al., 2006).

Chromosome pairing studies have shown that the A and A^t^ genomes are more closely related to each other than are the B and G genomes with A and A^t^ homoeologues pairing approximately 70% of the time compared to B-G homoeologues which pair approximately only 30% of the time (Feldman 1966). The ease with which the genetic variation in a wild species can be accessed is partly dependent on how closely related the genome(s) of the wild relative is to that of wheat, as the closer the relationship the higher the levels of homoeologous recombination possible. Hence, the pairing studies above would suggest that when crossing *T. timopheevii* to wheat, introgression of the A^t^ genome into the A genome is more likely to occur as compared to introgression of the G genome into the B genome.

A comprehensive set of molecular markers, polymorphic between wheat and *T. timopheevii*, is essential both during the generation of introgressions (for identification and characterisation) and also for the use of the introgressions in a breeding programme (in order to track the introgressions through the different generations). We earlier reported the generation of *T. timopheevii* introgression lines (Devi et al., 2019) with introgression identification and characterisation based on the Axiom^®^ Wheat Relative Genotyping Array (King et al., 2017). The array was extremely valuable for the initial screening of a large number of putative introgression lines but did not provide the flexibility required for screening different generations. A set of wheat chromosome-specific KASP markers was therefore generated for use on wheat/wild relative introgressions produced from 10 different wild relatives including *T. timopheevii* (Grewal et al., 2020a). These KASP markers were used to characterise 91 lines of wheat carrying homozygous introgressions from *T. timopheevii*. However, with the close relationship between the genomes of *T. timopheevii* and wheat, a relatively high level of recombination was expected, with the possibility of numerous smaller introgressions. Further KASP markers, polymorphic between *T. timopheevii* and bread wheat, have therefore been generated to provide a greater density of markers and to increase the detection power of the marker system.

## 2 Materials and Methods

### 2.1 Plant Material

Advanced back-crossed populations of wheat-*T. timopheevii* introgression lines, previously generated by Devi et al., (2019), were used as the starting material to generate homozygous ILs. Devi et al., (2019) crossed hexaploid wheat cv. Paragon *ph1/ph1* mutant (2n = 2x = 14) (obtained from the Germplasm Resource Unit (GRU) at the John Innes Centre) with *T. timopheevii*, (accession P95-99.1-1, obtained from the United States Department of Agriculture, USDA; 2n = 4x = 28), to produce F_1_ hybrids which were backcrossed with Paragon to generate BC_1_ populations. The BC_1_ individuals were recurrently backcrossed to produce BC_2_, BC_3_ and BC_4_ populations and self-fertilised at each generation.

Seeds were germinated in soil in seedling trays. Leaf tissues (1.5-inch leaf segment cut into pieces) were harvested from 3-week-old plants, immediately frozen on liquid nitrogen and stored at −80°C until nucleic acid extraction. Seedlings were vernalised for 4 weeks at 4°C after which the plants were transferred to 2L pots containing John Innes No. 2 soil, irrigated and fertilized using a drip irrigation system and maintained at 18–25°C under 16 h light and 8 h dark conditions. Plants were either self-fertilised or hand pollinated and crossed with the recurrent parent, wheat cv. Paragon, as the male parent.

#### 2.1.1 Breeding Scheme

The breeding scheme used to develop the homozygous wheat-*T. timopheevii* ILs is shown in Figure 1. A set of advanced backcrossed families were obtained through a previous study (Devi et al., 2019) and genotyped with KASP markers. Plants from different backcross families, carrying overlapping *T. timopheevii* segments from each of the 14 linkage groups were selected for further backcrossing and self-fertilisation. In the next generation, six to eight progenies from each selected plant were genotyped and those containing heterozygous segments of interest but still carrying two or more other (heterozygous or homozygous) *T. timopheevii* segments were selected for further backcrossing and self-fertilisation. Plants with potentially one or two homozygous *T. timopheevii* introgressions were self-fertilised and selected as homozygous introgression line (ILs). This process of plant selection through genotyping of segregating backcross and self-fertilised families was repeated for another generation of plants. At this point any plants showing one or more homozygous *T. timopheevii* segments were selected as homozygous ILs.

**Figure 1.**
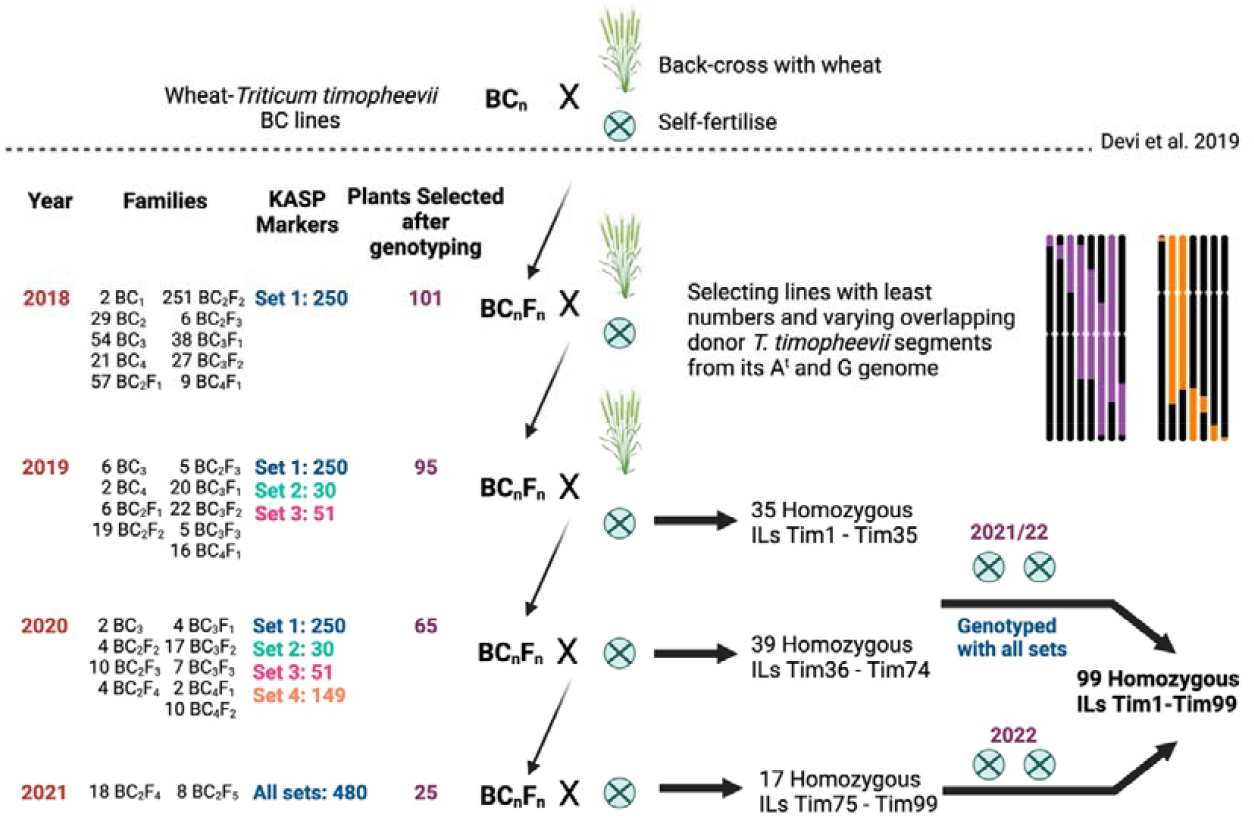
Breeding scheme used to generate homozygous wheat-*Triticum timopheevii* introgression lines (ILs) from advanced backcrossed lines generated by Devi et al., (2019). For each year (one generation per year), the number of families, KASP marker sets assessed, selected plants, and type of cross are shown. The backcrosses with the recurrent parents are represented by an × and a wheat plant, and the self-fertilisation rounds by an encircled ×. In each year a set of homozygous ILs were selected for additional rounds of self-fertilisation to create a panel of plants carrying overlapping *T. timopheevii* segments of varying sizes.

### 2.2 DNA Extraction

Leaf tissue samples were collected and freeze-dried in a deep-well plate and ground in a TissueLyser II (QIAGEN) for 4-6 minutes at a frequency of 25 Hz. Genomic DNA for sequencing and genotyping was extracted as described in Grewal et al., 2022.

Genomic DNA extraction for generation of probes for genomic *in situ* hybridisation analysis, was carried using the same protocol with an additional step of purification with phenol/chloroform at the end.

### 2.3 Development of KASP Markers

#### 2.3.1 Chromosome-specific SNP discovery

DNA was isolated from *T. timopheevii* accession P95-99.1-1, as described above, and a PCR-free library was prepared and sequenced on Illumina HiSeq 2500 to produce 250bp paired-end reads at a coverage of ~5x. SNP discovery followed a bioinformatics pipeline that produced a set of SNPs suitable for KASP assay design. In detail, the reads were mapped to the wheat cv. Chinese Spring reference genome assembly RefSeq v1.0 (International Wheat Genome Sequencing Consortium (IWGSC) et al., 2018) using BWA MEM version 0.7.13 (Li 2013) with the -M flag. The resulting alignments were filtered using SAMtools v1.4 (Li et al., 2009) to remove unmapped reads, supplementary alignments, improperly paired reads, and non-uniquely mapping reads (q<10). PCR duplicates were removed using Picard’s MarkDuplicates (DePristo et al., 2011). SNPs were called using BCFtools (Li, 2011) using the multiallelic model (-m). To remove SNPs unsuitable for KASP assays, we retained homozygous SNPs with a read depth greater than 10 and 100% mapped reads supporting the alternative allele and each position 50bp up and downstream of the SNP site having no INDELs, containing greater than 5 reads and >=80% of reads supporting the reference allele. This aims to remove sites with nearby variants that could disrupt the KASP assay. To prevent amplification of off-target (not the target wheat chromosome) regions in the genome, the remaining SNP sites, along with 50bp up and downstream were aligned to RefSeq v1.0 using BLASTn (Camacho et al., 2009). SNP sites were removed if in addition to the expected self-hit they produced an off-target hit with >=97% identity.

#### 2.3.2 KASP Assay Design

To design a KASP™ assay, the flanking sequence of a SNP was fed through the PolyMarker application (Ramirez-Gonzalez et al., 2015) which aligned the query sequence to wheat cv. Chinese Spring RefSeq v1.0 and provided two allele-specific primers and one common primer for each assay. A value of 1 in the *‘total_contigs’* column of the output *Primers* file validated the query SNP to be in a unique sequence region in the wheat genome RefSeq v1.0 assembly. The position of the KASP markers on wheat chromosomes in shown in Figure 2 and was generated through RIdeogram (Hao et al., 2020)

**Figure 2.**
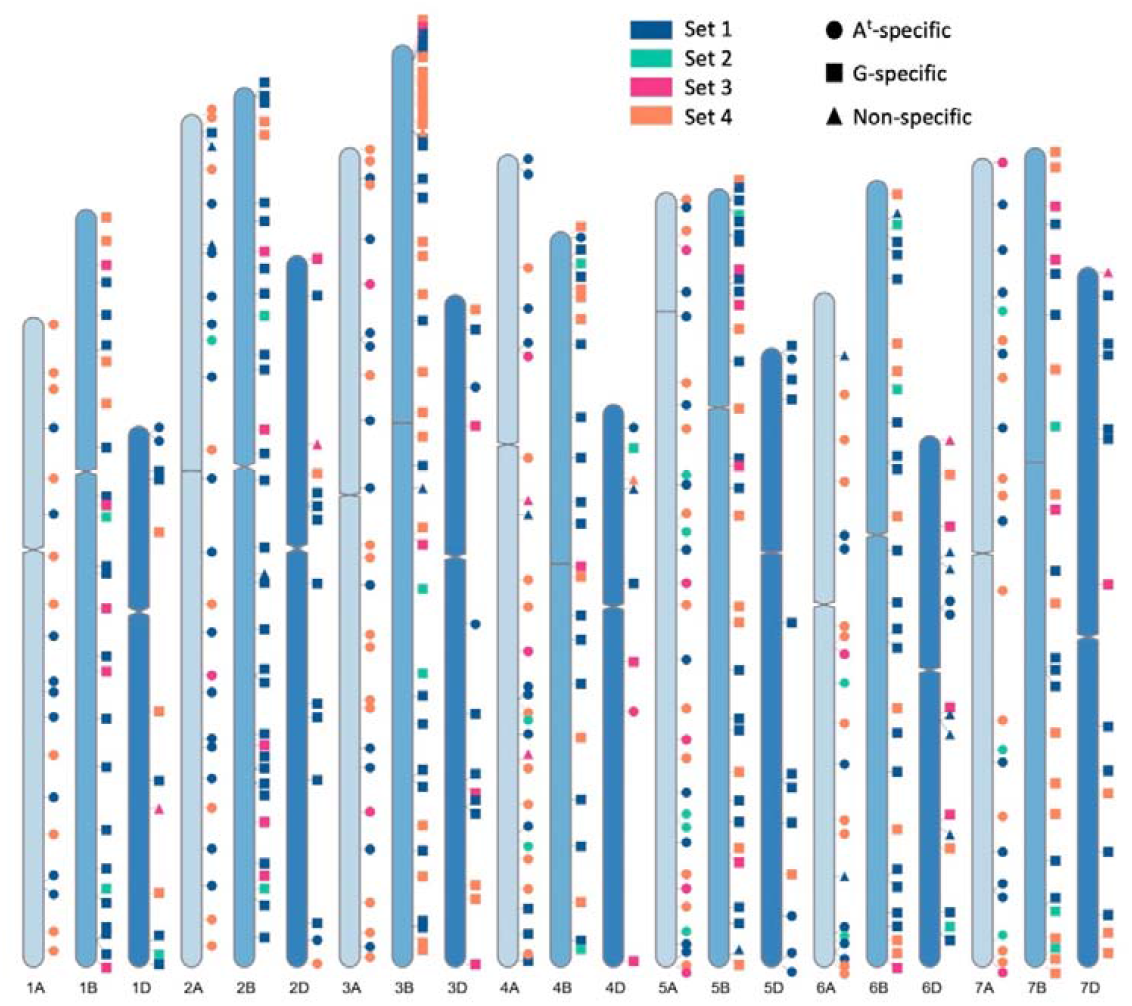
A graphical representation of physical positions of the four sets of chromosome-specific KASP markers, across all wheat chromosomes, used in this study to characterise wheat-*T. timopheevii* introgression lines. Sets 1, 2 and 3 were developed in previous works (Grewal et al., 2022; Grewal et al., 2021; Grewal et al., 2020a) and Set 4 was developed in this study to fill in the gaps between markers from the previous sets. Circles depict KASP markers that specifically detect A^t^ subgenome segments, squares depict markers that detect G subgenome segments and triangles represent markers that detect both subgenomes.

#### 2.3.3 Genotyping with KASP Markers

For genotyping during the breeding pipeline, four sets of KASP markers were used. The first three were developed during previous studies (Grewal et al., 2022; Grewal et al., 2021; Grewal et al., 2020a) and the fourth was developed during this work using chromosome-specific SNPs between wheat and *T. timopheevii*, discovered as described above.

Set 1 consisted of 250 chromosome-specific KASP markers previously reported to be polymorphic between wheat and *T. timopheevii* (codes between WRC0001-1000; Grewal et al., 2020a). KASP assays in Set 2 and 3 were originally designed to be polymorphic between wheat and other species and needed to be tested on *T. timopheevii* first to find those that were polymorphic between wheat and *T. timopheevii*. A subset of the KASP assays originally tested in those studies were selected for Set 2 and 3: those that did not fail at the PCR stage. Set 2 consisted of KASP assays designed to be tested on doubled haploid wheat-*T. urartu* ILs (codes between WRC1080-1308 and WRC1317-1393; Grewal et al., 2021) and Set 3 consisted of KASP assays designed to be tested on doubled haploid *wheat-Amblyopyrum muticum* ILs (codes between WRC1309-1316, WRC1394-1713, WRC1723-1872, WRC1894-1913, WRC1954-2113 and WRC2130-2169; Grewal et al., 2022). Set 4 consisted of the new KASP markers that were designed in this study (codes between WRC2170-2341 and WRC2366-2401).

The genotyping procedure was as described by Grewal et al., (2020b). Briefly, the genotyping reactions were set up using the automated PIPETMAX^®^ 268 (Gilson, UK) and performed in a ProFlex PCR system (Applied Biosystems by Life Technology) in a final volume of 5 μl with 1 ng genomic DNA, 2.5 μl KASP reaction mix (ROX), 0.068 μl primer mix and 2.43 μl nuclease free water. PCR conditions were set as 15 min at 94°C; 10 touchdown cycles of 10 s at 94°C, 1 min at 65–57°C (dropping 0.8°C per cycle); and 35 cycles of 15 s at 94°C and 1 min at 57°C. Fluorescence detection of the reactions was performed using a QuantStudio 5 (Applied Biosystems) and the data analysed using the QuantStudio™ Design and Analysis Software V1.5.0 (Applied Biosystems).

ChromoMap (Anand and Rodriguez Lopez 2022) was used to annotate each unique *T. timopheevii* segment introgressed into wheat and RIdeogram (Hao et al., 2020) was used to annotate the KASP marker positions on the wheat genome.

### 2.4 *In situ* Hybridisation

Preparation of the root-tip metaphase chromosome spreads for *in situ* hybridisation analysis was as described in King et al., (2017) and Grewal et al., (2017) with a few modifications. Briefly, two roots (2 cm) from a 2-day old seedling were excised and treated with nitrous oxide gas at 10 bar for 2 h. Treated roots were fixed in 90% acetic acid for 15 min and then washed three times with water, on ice. The root tip was dissected and digested in 20 μL of 1% pectolyase Y23 and 2% cellulase Onozuka R-10 (Yakult Pharmaceutical, Tokyo) solution for 50 min at 37 °C and then washed three times with 70% ethanol. The root tips were crushed in 70% ethanol, and the cells collected by centrifugation at 3000 x g for 1 min, briefly dried and then re-suspended in 20–30 μL of 100% acetic acid before being placed on ice. The cell suspension was dropped onto glass slides (6–7 μL per slide) in a moist box and dried slowly under cover. Slide denaturation was performed at 80 °C for 5 min and probes were hybridized at 55 °C for 16 h. Preparation of probes for genomic and fluorescent *in situ* hybridisation was as described below.

Post-hybridization, all slides were washed with 2 × saline-sodium citrate (SSC) buffer for 1 min and counter-stained with DAPI (4’,6-diamidino-2-phenylindole). Metaphases were detected using a high-throughput, fully automated Zeiss Axio ImagerZ2 upright epifluorescence microscope (Carl Zeiss Ltd., Germany). Image capture was performed using a MetaSystems Coolcube 1m CCD camera and image analysis was carried out using Metafer4 (automated metaphase image capture) and ISIS (image processing) software (Metasystems GmbH, Germany).

#### 2.4.1 Multi-colour Genomic *In situ* Hybridisation (mcGISH)

Genomic DNA from *T. urartu* (to detect the A-genome), *Ae. speltoides* (to detect the B-genome), and *Aegilops tauschii* (to detect the D-genome) were isolated as described above and labelled to be used as probes. Two micrograms of genomic DNA was labelled by nick translation (Rigby et al., 1977) as follows: (1) *T. urartu* with ChromaTide™ Alexa Fluor™ 488-5-dUTP (Invitrogen; C11397; coloured green), (2) *Ae. speltoides* with DEAC-dUTP (Jena Bioscience; NU-803-DEAC; coloured blueish purple), and (3) *Ae. tauschii* with ChromaTide™ Alexa Fluor™ 594-5-dUTP (Invitrogen; C11400; coloured red). Slides were probed using 150 ng of *T. urartu*, 150 ng of *Ae. speltoides* and 300 ng of *Ae. tauschii*, in the ratio 1:1:2 (green: blue: red). No blocking DNA was used.

#### 2.4.1 Multi-colour Fluorescent *In situ* Hybridisation (mcFISH)

Mc-FISH was carried out as described by Grewal et al., (2018). The pSc119.2 clone (McIntyre et al., 1990), a 120-bp repetitive DNA sequence derived from rye (*Secale cereale* L.) and inserted into the plasmid pBR322, and the pAs.1 clone (Rayburn and Gill 1986), a 1kb repetitive DNA sequence isolated from *Ae. tauschii* and inserted into the pUC8 plasmid, were nick-labelled (Rigby et al., 1977) with Alexa Fluor 488-5-dUTP (green) and Alexa Fluor 594-5-dUTP (red), respectively. Classification of pSc119.2- and pAs.1-labelled chromosomes followed the nomenclature of Badaeva et al., 2016.

## 3 Results

### 3.1 Development of homozygous introgression lines (ILs)

Advanced backcrossed populations of wheat-*T. timopheevii* introgression lines developed by Devi et al., 2019, segregating for various *T. timopheevii* segments, were propagated through backcrossing and self-fertilisation and genotyped at every generation with chromosome-specific KASP markers (Grewal et al., 2020a; Grewal et al., 2021; Grewal et al., 2022) as shown in Figure 1. Plants carrying segments of various sizes and from different linkage groups were selected to create a panel of overlapping *T. timopheevii* introgressions from each of its linkage groups in the wheat background.

In 2018, a previously developed set of chromosome-specific KASP markers (Grewal et al., 2020a) consisting of 250 markers polymorphic between wheat and *T. timopheevii*, and designated as Set 1, were used to genotype 1685 advanced backcrossed lines (Devi et al., 2019) belonging to 494 BC_n_F_n_ families (Figure 1). The genotyping identified 101 plants that carried unique *T. timopheevii* segments but since these lines also had multiple segments or whole chromosomes from *T. timopheevii*, they were taken forward to another generation either through backcrossing with wheat cv. Paragon or through self-pollination.

The next generation, segregating for the desired *T. timopheevii* segments and consisting of 772 plants from 101 BC_n_F_n_ families, were genotyped with the Set 1 KASP markers in 2019. As shown in Figure 2, this set of markers had significant gaps on many chromosomes and thus, additional chromosome-specific KASP markers were sought to fill in those gaps. Simultaneously to this work, KASP markers were being developed to characterise doubled haploid wheat ILs from *T. urartu* and *Am. muticum* (Grewal et al., 2021; Grewal et al., 2022). These two groups of chromosome-specific KASP assays consisting of 224 markers developed for *T. urartu* and 425 markers developed for *Am. muticum* were tested on *T. timopheevii* and its derived segregating BC_n_F_n_ families. From the first group, 30 were found to be diagnostic for *T. timopheevii*, designated as Set 2, and from the second group 51 markers were validated for *T. timopheevii* and designated as Set 3 (Figure 2; Table 1). After genotyping with the three sets of KASP markers, 35 plants were selected as carrying one or two homozygous *T. timopheevii* segments and self-fertilised to create stable introgression lines (ILs). An additional 60 plants were selected as carrying segments of interest and were further backcrossed and self-fertilised either because the segment(s) were still heterozygous or to segregate multiple segments through a family to create ILs with fewer *T. timopheevii* segments.

**Table 1.** Number of chromosome-specific markers across the different sets of markers tested, average and maximum distance between markers and *T. timopheevii* subgenome-specific markers per wheat chromosome.

| Wheat Chromosome | Set 1 | Set 2 | Set 3 | Set 4 | Total | Average distance between markers (Mbp) | Max. distance between 2 markers (Mbp) | A<sup>t</sup>-specific | G-specific | Non-specific |
| --- | --- | --- | --- | --- | --- | --- | --- | --- | --- | --- |
| 1A | 9 | 0 | 0 | 10 | 19 | 29.68 | 46.30 | 19 | 0 | 0 |
| 1B | 16 | 2 | 5 | 4 | 27 | 24.62 | 57.59 | 0 | 27 | 0 |
| 1D | 7 | 1 | 1 | 3 | 12 | 38.11 | 164.34 | 2 | 9 | 1 |
| 2A | 17 | 1 | 1 | 8 | 27 | 27.89 | 67.47 | 24 | 1 | 2 |
| 2B | 25 | 2 | 5 | 2 | 34 | 22.89 | 62.09 | 0 | 33 | 1 |
| 2D | 10 | 0 | 2 | 2 | 14 | 43.46 | 136.34 | 2 | 11 | 1 |
| 3A | 11 | 0 | 2 | 13 | 26 | 27.81 | 61.95 | 26 | 0 | 0 |
| 3B | 17 | 2 | 2 | 22 | 43 | 18.85 | 75.71 | 0 | 41 | 2 |
| 3D | 7 | 0 | 3 | 3 | 13 | 43.97 | 181.60 | 2 | 11 | 0 |
| 4A | 12 | 2 | 4 | 10 | 28 | 25.68 | 99.25 | 22 | 3 | 3 |
| 4B | 14 | 2 | 1 | 7 | 24 | 26.94 | 66.91 | 1 | 23 | 0 |
| 4D | 3 | 1 | 3 | 1 | 8 | 56.65 | 228.68 | 2 | 4 | 2 |
| 5A | 11 | 5 | 5 | 11 | 32 | 21.51 | 60.90 | 32 | 0 | 0 |
| 5B | 18 | 1 | 4 | 9 | 32 | 21.61 | 83.29 | 0 | 31 | 1 |
| 5D | 11 | 0 | 0 | 1 | 12 | 43.54 | 204.45 | 5 | 7 | 0 |
| 6A | 8 | 2 | 1 | 10 | 21 | 28.09 | 70.81 | 19 | 0 | 2 |
| 6B | 17 | 2 | 1 | 8 | 28 | 24.86 | 61.64 | 0 | 27 | 1 |
| 6D | 9 | 1 | 4 | 2 | 16 | 27.86 | 97.57 | 2 | 8 | 6 |
| 7A | 10 | 3 | 2 | 9 | 24 | 29.47 | 118.70 | 24 | 0 | 0 |
| 7B | 9 | 3 | 3 | 11 | 26 | 27.80 | 61.86 | 0 | 26 | 0 |
| 7D | 9 | 0 | 2 | 3 | 14 | 42.58 | 133.02 | 0 | 13 | 1 |
| **Total** | **250** | **30** | **51** | **149** | **480** |  |  | **182** | **275** | **23** |

In 2020, the next generation consisting of 356 plants from 60 BC_n_F_n_ families (Figure 1) were genotyped with Set 1, 2 and 3 KASP markers. The aim was to have a KASP marker at every 60-70 Mbp across the A and B genomes of wheat for genotyping of these lines. However, as shown in Figure 2, there were still significant gaps between markers from Sets 1-3 on some A and B genome chromosomes.

To fill in these gaps, new KASP markers were developed through discovery of chromosome-specific SNPs between *T. timopheevii* and wheat. After aligning *T. timopheevii* sequence reads against the wheat Chinese Spring RefSeqv1.0 assembly (International Wheat Genome Sequencing Consortium (IWGSC) et al, 2018), variant calling and filtering for KASP assay suitable flanking regions, 36,001 SNPs were generated between wheat and *T. timopheevii*. SNPs with significant off-target hits were removed to give 6957 chromosome-specific SNPs (Supplemental data S1). Of these 208 SNPs were selected at positions that filled the most prominent gaps between markers in Sets 1-3, converted to KASP assays and tested on the segregating families. The results of genotyping showed that 39 assays failed at the PCR stage, 20 assays were not robust in separating heterozygotes and homozygotes and thus, 149 assays were able to successfully detect *T. timopheevii* segments in a wheat background. The latter were designated as Set 4 (Figure 2; Table 1).

Based on the genotyping results of markers from Sets 1-4, a further 39 ILs were selected as carrying only homozygous *T. timopheevii* segments (Figure 1). Along with the 35 ILs from the previous generation, all 74 ILs were self-fertilised for two further generations and genotyped with all 480 KASP markers from Sets 1-4. These lines were code named Tim1 to Tim74.

A further 26 plants were selected, from this last generation, that had a combination of heterozygous and homozygous desirable *T. timopheevii* segments. Their corresponding 26 BC_n_F_n_ families were taken forward through the germination of 89 plants. After genotyping with all 480 KASP markers, 25 were selected as homozygous for all introgressions and are currently undergoing the first of two rounds of self-fertilisation. These lines were code named Tim75 to Tim99 and will be undergo a further two rounds of multiplication.

### 3.2 Distribution and specificity of KASP markers

In total 480 chromosome-specific KASP markers were used to genotype the wheat-*T. timopheevii* ILs (Table 1; Figure 2; Supplemental data S2). These markers were designed to be well-distributed across the wheat chromosomes. The average distance between markers ranged from 21.51 – 29.68 Mbp on the A genome chromosomes, 18.85 – 27.80 Mbp on the B genome chromosomes and 27.86 – 56.65 Mbp on the D genome chromosomes (Table 1). The latter had the largest distances between two markers but there were also some significant gaps between markers on chromosomes 4A, 5B and 7A (Table 1; Figure 2).

In addition to being able to only detect one wheat chromosome, the KASP markers developed for *T. timopheevii* were, in the majority of cases, also able to specifically detect either the A^t^ or the G subgenome chromosomes of *T. timopheevii*. Of the 480 KASP markers used to genotype the ILs, 182 were found to detect only A^t^ subgenome segments, 275 detected G subgenome segments and 23 were non-specific for a *T. timopheevii* subgenome (Table 1, Figure 2, Supplemental Data S2).

### 3.3 Introgressions from *T. timopheevii* in wheat

A range of different sized segments from each *T. timopheevii* chromosome of the A^t^ and G subgenomes were found present in the ILs and are annotated in Figures 3 and 4, respectively. In total, there were 89 unique segments from the A^t^ subgenome of *T. timopheevii* and a whole chromosome 4A^t^ (Figure 3). It was estimated that all A^t^ chromosomes, except chromosome 3A^t^, were introgressed entirely into wheat through overlapping segments with approximately 93.4% of chromosome 3A^t^ also being transferred. Figure 4 shows the 74 unique segments from the G subgenome of *T. timopheevii* that were introgressed into the ILs. It was estimated that 92.4% and 98.3% of chromosomes 4G and 6G, respectively, were introgressed in overlapping segments in addition to the entirety of the rest of the G subgenome chromosomes. The most segments introgressed into wheat were from chromosome 5A^t^ (25) and chromosome 2G (20) of *T. timopheevii*. Together, approximately 98.9% of the *T. timopheevii* genome had been introgressed into bread wheat mainly via small and large overlapping segments. The full set of chromosome-specific KASP markers detected 183 introgressions from the A^t^ subgenome and 126 from the G subgenome of *T. timopheevii* in the 99 ILs developed in this work.

**Figure 3.**
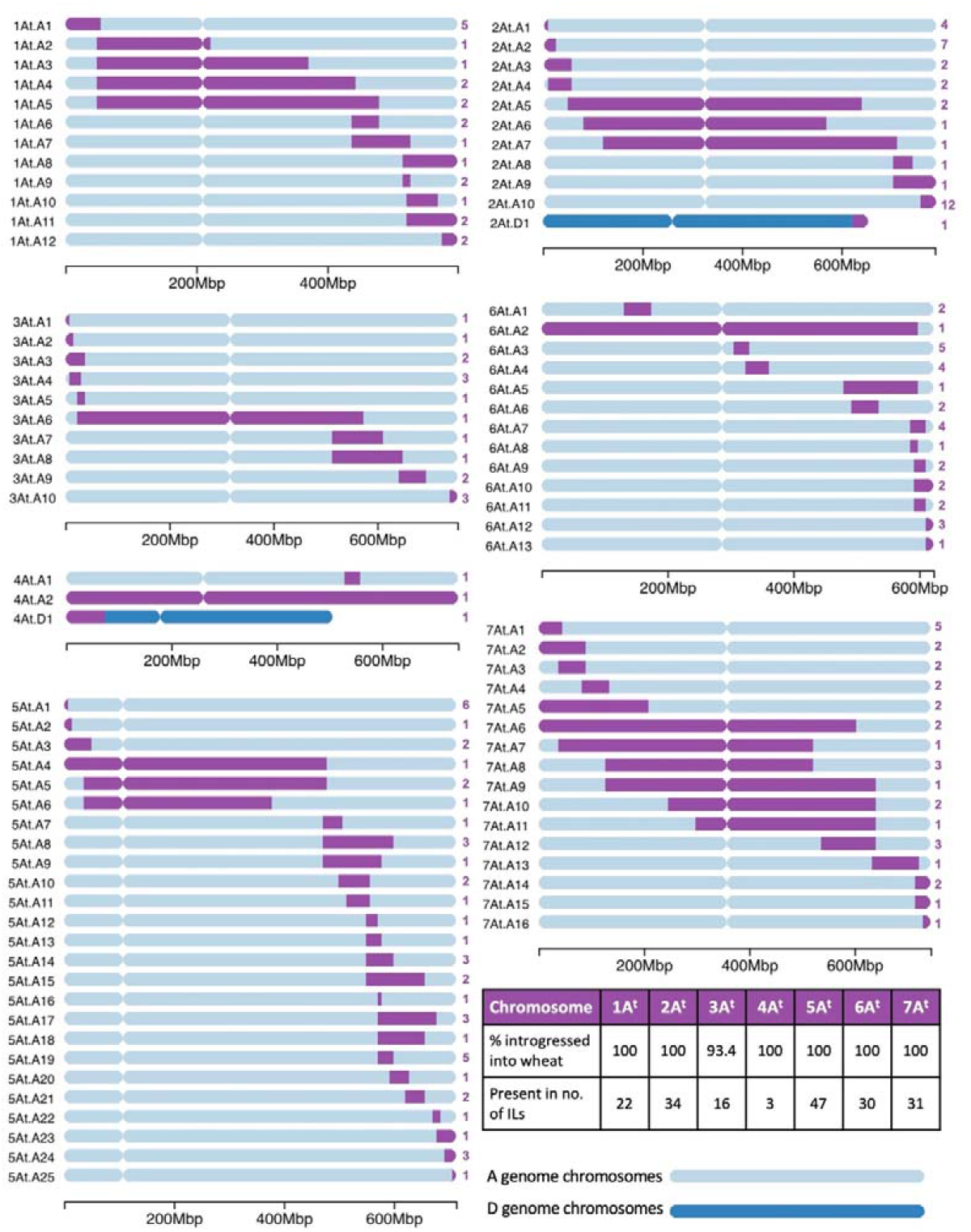
A graphical representation of the size and location of unique introgressions from the A^t^ genome of *T. timopheevii* (purple) in wheat A and D genome chromosomes (shades of blue) for each of the seven linkage groups. On the left of each ideogram is the label of each unique introgression and on the right is the number (in purple) of introgression lines (ILs) the segment is found in. A table on the bottom right shows the percentage of each A^t^ chromosome that has been introgressed into the wheat genome and the number of ILs that contains a homozygous segment from each of the A^t^ chromosomes. A scale at the bottom of each linkage group shows the size of the ideograms.

**Figure 4.**
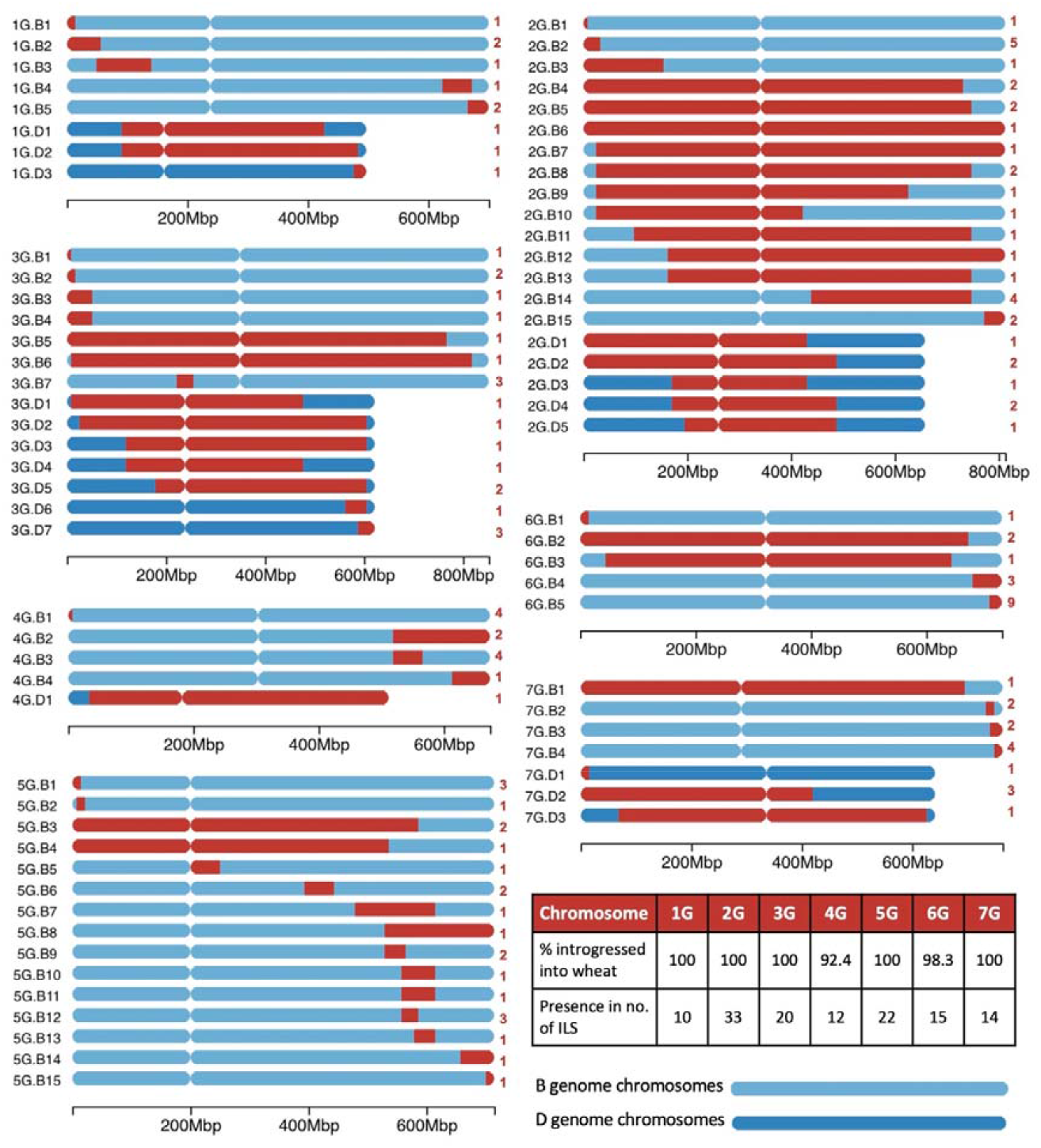
A graphical representation of the size and location of unique introgressions from the G genome of *T. timopheevii* (red) in wheat B and D genome chromosomes (shades of blue) for each of the seven linkage groups. On the left of each ideogram is the label of each unique introgression and on the right is the number (in red) of introgression lines (ILs) the segment is found in. A table on the bottom right shows the percentage of each G chromosome that has been introgressed into the wheat genome and the number of ILs that contains a homozygous segment from each of the G chromosomes. A scale at the bottom of each linkage group shows the size of the ideograms.

Table 2 details which of these segments are carried by each of the ILs Tim1-99 and a diagnostic KASP marker for each segment is provided in Supplemental Data S3. Genotyping of ILs Tim1 to 35 with Set 4 KASP markers identified additional homozygous *T. timopheevii* segments that had not been previously identified by markers in Sets 1-3. It was also found that ILs Tim14 and Tim29 had the same genotype according to the markers used. Each IL contained between one and eight homozygous introgressions from *T. timopheevii* with an average of 3 segments per IL (Table 2).

**Table 2.** Number and description of *T. timopheevii* introgressions present in each wheat-*T. timopheevii* introgression line (IL), corresponding to the segments depicted in Figures 3 and 4).

| IL Name | No. of Introgressions | Introgressions present (as shown in Figures 3 and 4) | IL Name | No. of Introgressions | Introgressions present (as shown in Figures 3 and 4) |
| --- | --- | --- | --- | --- | --- |
| Tim1 | 3 | 2At.A5, 6At.A9, 7G.B4 | Tim51 | 4 | 2G.B5, 3At.A3, 5At.A23, 7At.A10 |
| Tim2 | 3 | 1At.A9, 3G.B2, 7G.B3 | Tim52 | 2 | 3At.A3, 7At.A10 |
| Tim3 | 3 | 2At.A10, 5At.A17, 7At.A8 | Tim53 | 2 | 1At.A2, 2G.B14 |
| Tim4 | 4 | 5At.A1, 5At.A8, 6G.B4, 7At.A1 | Tim54 | 1 | 2G.B14 |
| Tim5 | 3 | 1At.A12, 2G.D2, 3G.B4 | Tim55 | 7 | 1At.A1, 2G.B10, 3At.A10, 3G.D1, 5At.A12, 6At.A12, 6G.B5 |
| Tim6 | 3 | 2G.D4, 3G.B5, 6G.B5 | Tim56 | 5 | 1At.A4, 2G.B14, 3G.D2, 5At.A24, 7At.A5 |
| Tim7 | 2 | 2At.A10, 7G.D2 | Tim57 | 5 | 1At.A1, 2At.A10, 3At.A10, 7At.A1, 7At.A12 |
| Tim8 | 2 | 2At.A10, 5G.B3 | Tim58 | 2 | 1At.A10, 5At.A16 |
| Tim9 | 4 | 2At.A10, 5G.B3, 7At.A9, 7G.B4 | Tim59 | 2 | 3G.D7, 4G.B3 |
| Tim10 | 5 | 2At.A10, 2G.B15, 3At.A1, 5At.A13, 5G.B14 | Tim60 | 1 | 4G.B3 |
| Tim11 | 2 | 2G.D4, 6At.A3 | Tim61 | 2 | 1G.D2, 6At.A2 |
| Tim12 | 3 | 2G.D5, 6At.A3, 7G.D2 | Tim62 | 1 | 2G.B11 |
| Tim13 | 3 | 7At.A1, 7At.A12, 7G.D2 | Tim63 | 1 | 2G.D3 |
| Tim14/29 | 3 | 1At.A5, 2At.A1, 4G.B1 | Tim64 | 1 | 5At.A11 |
| Tim15 | 2 | 1At.A7, 1G.B1 | Tim65 | 1 | 3G.D6 |
| Tim16 | 2 | 2At.A10, 3G.D5 | Tim66 | 1 | 5G.B10 |
| Tim17 | 2 | 4G.B2, 5At.A24 | Tim67 | 1 | 7At.A3 |
| Tim18 | 3 | 5G.B15, 6At.A4, 6G.B3 | Tim68 | 2 | 4G.B3, 7At.A3 |
| Tim19 | 3 | 1G.D1, 4G.B1, 6At.A1 | Tim69 | 2 | 4G.B4, 6At.A4 |
| Tim20 | 1 | 2G.B7 | Tim70 | 2 | 2G.B2, 6At.A9 |
| Tim21 | 2 | 2At.A1, 4G.B3 | Tim71 | 2 | 2G.B2, 6At.A7 |
| Tim22 | 1 | 6At.A7 | Tim72 | 1 | 2G.B2 |
| Tim23 | 1 | 5At.A10 | Tim73 | 6 | 2At.A3, 2G.B3, 2G.B15, 3G.D3, 5At.A19, 6G.B5 |
| Tim24 | 2 | 3G.B1, 7G.B1 | Tim74 | 4 | 1At.A1, 1At.A12, 2G.B13, 7At.A6 |

| **IL Name** | **No. of Introgressions** | **Introgressions present (as shown in Figures 3 and 4)** | **IL Name** | **No. of Introgressions** | **Introgressions present (as shown in Figures 3 and 4)** |
| --- | --- | --- | --- | --- | --- |
| Tim25 | 2 | 2G.D1, 5At.A10 | Tim75 | 4 | 2G.B2, 6At.A7, 7G.B4, 7G.D3 |
| Tim26 | 1 | 2G.B12 | Tim76 | 3 | 2G.B2, 4At.A1, 6G.B2 |
| Tim27 | 4 | 2G.B9, 3G.B7, 5At.A20, 6G.B5 | Tim77 | 5 | 1At.A1, 2At.A2, 3At.A8, 5At.A2, 5At.A25 |
| Tim28 | 3 | 1At.A1, 5G.B8, 6At.A3 | Tim78 | 3 | 1G.D3, 2At.A4, 6G.B2 |
| Tim30 | 1 | 2At.A5 | Tim79 | 7 | 1G.B5, 2At.A2, 2At.A7, 5At.A1, 6G.B1, 7At.A4, 7At.A13 |
| Tim31 | 3 | 5At.A3, 5At.A8, 6G.B4 | Tim80 | 4 | 1G.B5, 2At.A2, 6At.A5, 7At.A4 |
| Tim32 | 5 | 2At.A6, 3At.A10, 5At.A18, 6At.A4, 7At.A1 | Tim81 | 3 | 5At.A19, 5G.B11, 6At.A10 |
| Tim33 | 2 | 2G.B14, 7At.A5 | Tim82 | 5 | 3At.A2, 3G.B6, 5At.A24, 6At.A13, 6G.B5 |
| Tim34 | 3 | 2At.A3, 2At.A10, 3G.D5 | Tim83 | 2 | 2G.B1, 5G.B7 |
| Tim35 | 3 | 1At.A9, 3G.B2, 7G.B3 | Tim84 | 4 | 3G.B7, 4At.D1, 6G.B5, 7G.B2 |
| Tim36 | 2 | 2G.B5, 4G.B1 | Tim85 | 3 | 5G.B5, 6G.B5, 7G.B2 |
| Tim37 | 4 | 1At.A4, 4G.B2, 5At.A4, 7At.A6 | Tim86 | 4 | 1At.A6, 2At.D1, 5At.A17, 7At.A8 |
| Tim38 | 3 | 1At.A6, 5At.A17, 7At.A8 | Tim87 | 2 | 3G.B3, 4At |
| Tim39 | 3 | 2At.A10, 2G.B4, 5At.A19 | Tim88 | 6 | 2At.A4, 3At.A9, 5At.A14, 5G.B1, 5G.B12, 6At.A10 |
| Tim40 | 3 | 2At.A10, 2G.B4, 3At.A4 | Tim89 | 6 | 2At.A2, 3At.A6, 3At.A9, 5At.A14, 5G.B1, 6At.A11 |
| Tim41 | 2 | 2At.A10, 7At.A1 | Tim90 | 7 | 2At.A2, 3At.A5, 3At.A7, 5At.A14, 5G.B1, 5G.B13, 6At.A11 |
| Tim42 | 6 | 1At.A8, 5At.A1, 5At.A9, 6At.A8, 6At.A12, 7G.D1 | Tim91 | 6 | 1G.B3, 5At.A3, 5At.A8, 6At.A2, 6G.B5, 7At.A15 |
| Tim43 | 4 | 1G.B4, 2G.B8, 5At.A1, 5At.A7 | Tim92 | 1 | 7At.A9 |
| Tim44 | 2 | 5At.A6, 6At.A7 | Tim93 | 7 | 2At.A2, 2G.B6, 3G.D4, 5At.A19, 6G.B5, 7At.A16, 7G.B4 |
| Tim45 | 4 | 1G.B2, 5At.A5, 5At.A15, 6At.A3 | Tim94 | 8 | 2At.A1, 2At.A10, 4G.D1, 5G.B6, 5G.B12, 6At.A4, 7At.A2, 7At.A14 |
| Tim46 | 6 | 1G.B2, 2G.D2, 5At.A5, 5At.A15, 6At.A3, 6At.A12 | Tim95 | 5 | 5At.A22, 5G.B6, 5G.B12, 7At.A2, 7At.A14 |
| Tim47 | 3 | 3G.D7, 5G.B2, 7At.A7 | Tim96 | 2 | 1At.A3, 5At.A21 |
| Tim48 | 2 | 3G.D7, 5G.B4 | Tim97 | 7 | 1At.A11, 2G.B8, 3At.A4, 5G.B9, 6At.A6, 6G.B4, 7At.A11 |
| Tim49 | 2 | 2At.A2, 2At.A9 | Tim98 | 5 | 1At.A11, 3At.A4, 5At.A1, 5G.B9, 6At.A6 |
| Tim50 | 1 | 5At.A21 | Tim99 | 3 | 2At.A8, 3G.B7, 5At.A19 |

The wheat chromosome-specific KASP markers also showed that the majority of the A^t^ subgenome segments were introgressed into the A genome chromosomes of wheat (At.A; Figure 3) and the G subgenome segments into the B genome of wheat (G.B; Figure 4). However, only two A^t^ segments were found to have recombined with the D genome of wheat (At.D; Figure 3) compared to 19 segments from the G subgenome (G.D; Figure 4). Multi-colour GISH (mcGISH) was used to validate these D genome introgressions. Figure 5A shows mcGISH images of introgressions 4At.D1 and 2At.D1 where small A^t^ subgenome segments from *T. timopheevii* chromosomes 4A^t^ and 2A^t^ (shown in green) had recombined with chromosomes 4D and 2D of wheat (shown in red), respectively. Ideograms representing the KASP marker genotyping indicating the introgressed segments in orange (Figure 5B) matched the corresponding recombination breakpoints as indicated with white arrows in Figure 5A. Of the 19 G.D introgressions, mcGISH images of 6 introgressions, 3G.D1-D3 and 3G.D5-D7, each containing a different sized segments from chromosome 3G of *T. timopheevii* (shown in blue) recombined with chromosome 3D of wheat (shown in red) are shown in Figure 5C. Ideograms representing the KASP marker genotyping indicating each of the introgressed segments in orange are shown below in Figure 5D which correspond to the recombination breakpoints of the introgressions as indicated by white arrows in Figure 5C. It is not possible to detect At.A and G.B introgressions with mcGISH and thus, mc-FISH was used to validate the KASP marker genotyping by analysing three introgressions, 1At.A4, 2G.B3 and 3G.B3, which were large enough to span a diagnostic FISH signal for *T. timopheevii*. Images in Figures 5E, 5F and 5G show the FISH karyotypes for the indicated chromosomes in wheat cv. ‘Chinese Spring’, *T. timopheevii* (accession P95-99.1-1) and the wheat-*T. timopheevii* ILs, respectively. Figure 5H represents the KASP genotyping of those introgressions in each IL. The gain of the *T. timopheevii* signal (as indicated by the white arrow) in Tim14 validates the introgression of chromosome 1A^t^ in chromosome 1A of wheat which otherwise lacks this signal. The loss of the wheat signal on chromosome 2BS (as indicated by the white arrow) in Tim73 indicates the presence of the 2G segment from *T. timopheevii* which otherwise lacks this signal on chromosome 2GS. In Tim73, the gain of the *T. timopheevii* signal from chromosome 3G validates the 3G.D3 introgression in chromosome 3D which otherwise lacks this signal on 3DL.

**Figure 5.**
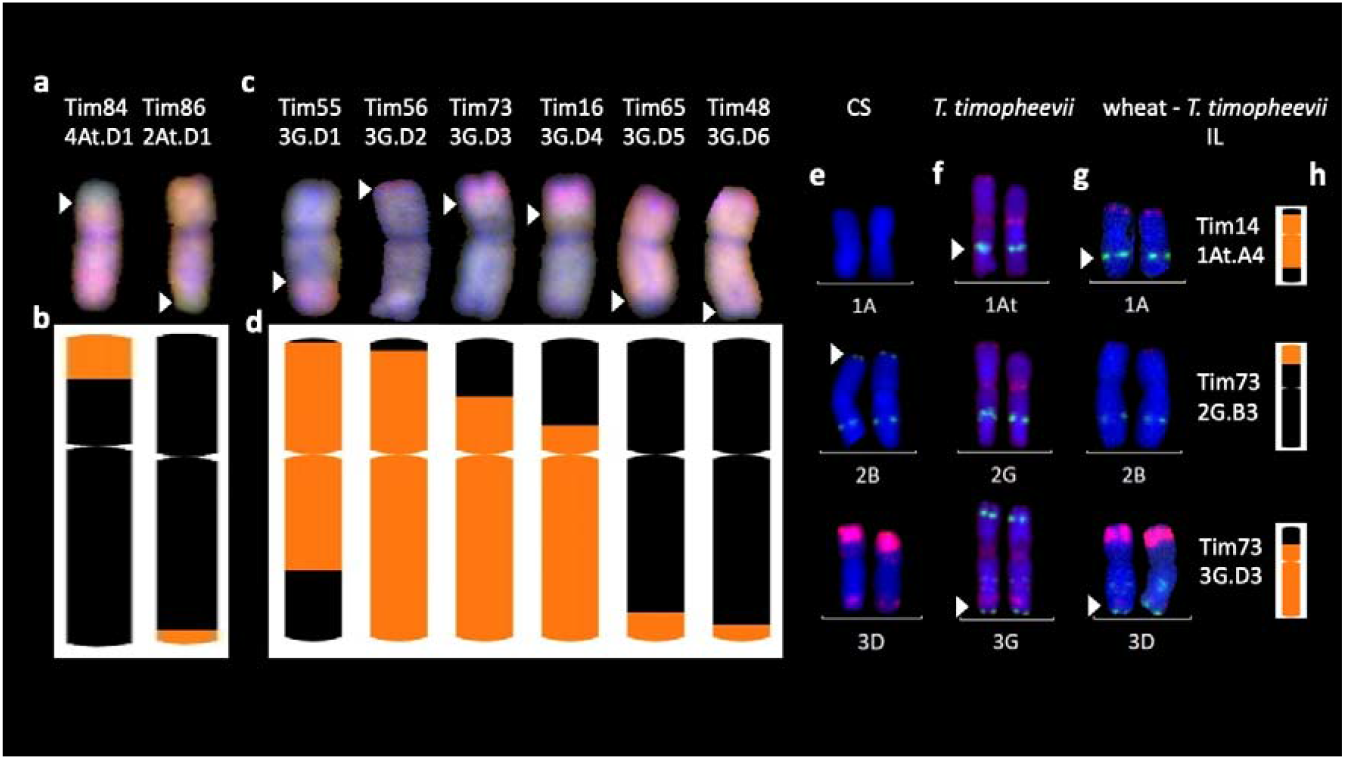
Multi-colour *in situ* hybridisation analysis of root metaphase spreads of chromosomes from wheat-*T. timopheevii* introgression lines (ILs). (A, C) multi-colour genomic *in situ* hybridisation (mcGISH) of ILs with At.D and G.B introgressions, respectively. Green probe detects the A^t^ subgenome and blue detects the G subgenome of *T. timopheevii*. Red detects the D genome of wheat. (E-G) multi-colour fluorescence *in situ* hybridisation (mcFISH) of (E) wheat cv. Chinese Spring (CS), (F) T. timopheevii and (G) ILs where the green and red signals show pSc119.2 and pAs.1 binding sites, respectively. Images for (E) and (F) were taken from Devi et al., 2019. (B, D, H) a graphical representation of the introgressions present in these lines obtained through chromosome-specific KASP marker analysis depicting ideograms of the corresponding ILs shown either above with mcGISH (A, C) or on the left with mcFISH (E-G). Size of introgressions from A^t^ and G subgenomes is indicated in orange in a wheat chromosome shown in black. White arrows indicate the corresponding recombination breakpoints in the mcGISH and mcFISH images.

## 4 Discussion

The main aims of this work were (i) the development of a set of chromosome-specific KASP markers, well spread across the A and B genomes of wheat, for the genotyping of the homozygous lines and (ii) to generate homozygous introgression lines (ILs) from the advanced backcrossed lines generated by Devi et al., (2019), thus enabling the multiplication and distribution of the lines for trait phenotyping.

The selection of plants with segments from *T. timopheevii* of various sizes and from different linkage groups was enabled through the use of chromosome-specific KASP markers. However, the markers needed to be of a sufficient density across all linkage groups in order to detect all introgressions present (including very small segments) and to accurately assess the size of a segment. The detection of small segments is particularly important in wheat-*T. timopheevii* ILs as the two subgenomes of *T. timopheevii* share considerable similarities to two of the genomes of wheat having originated from the same progenitors, *T. urartu* (A and A^t^ genomes) (Dvorak et al., 1993) and an *Ae. speltoides*-like species (B and G genomes) (Dvorak and Zhang 1990). The more closely related two genomes were, the higher the level of recombination is possible between the two.

### 4.1 Development of chromosome-specific KASP markers

Prior to this work other studies have developed molecular marker resources for the detection of *T. timopheevii* segments in a wheat background (Ji et al., 2007; Uhrin et al., 2012; Timonova et al., 2013; Devi et al., 2019) but none have reported the density and distribution across the wheat genome that has been achieved in this work. The aim in this study was to have a KASP markers at every 60-70 Mbp across the A and B genomes of wheat in order to carry out accurate genotyping of the ILs. Sets 1, 2 and 3 were designed either for multi-species use (Set 1) or were designed with the target of filling gaps between wheat and other wild relatives using SNPs discovered in those species (Set 2-*T. urartu* and Set 3 – *Am. muticum*). As can be seen in Figure 2, most of the chromosomes still had significant gaps between the markers of these 3 sets. The KASP markers of Set 4, however, were designed to target the gaps based on SNPs between wheat and *T. timopheevii*. The strategy of custom filtering to remove SNPs with significant off-target hits and only retain those suitable for KASP assay design resulted in 72% of the chromosome-specific KASP assays designed successfully able to detect the presence of *T. timopheevii* segments introgressed into wheat chromosomes. It is difficult to compare the success of this strategy with some of those used previously as the early panels of KASP assays were mainly designed to be polymorphic between wheat and more than one wild relative. However, the same strategy was recently followed to increase the number of SNPs obtained that were polymorphic between wheat and *Am. muticum* (Grewal et al., 2022). In that work only 48% of the KASP assays designed functioned to detect *Am. muticum* DNA in a wheat background. The difference in the success rates might have been due to *Am. muticum* being an outbreeder and thus highly heterozygous while *T. timopheevii* is an inbreeder with a much lower level of heterozygosity resulting in robust KASP assay primers with fewer SNPs in the flanking regions of the SNP.

Combined, the 4 sets of markers now give excellent genome coverage with only 3 relatively significant gaps still needing to be filled (1 each on 4A, 5B and 7A) with the largest of the gaps on 7A (118.7 Mbp; Table 1). The new, higher density of markers identified new homozygous segments in ILs Tim1 to 35 which had previously been genotyped with KASP marker sets 1, 2 and 3, highlighting the importance of increased marker density. The extra KASP markers will also enable these newly identified introgressions to be tracked in breeding programmes. A high density of markers at the end of chromosome 3B short arm can also be seen in Figure 2. This region was specifically targeted as recent phenotyping has shown this region of chromosome 3G to be associated with resistance to Fusarium Head Blight. The higher density of SNPs in this region demonstrates that the strategy of SNP discovery and KASP development employed will be successful for filling the remaining gaps. It is possible that even with the new, higher density of KASP markers, small introgressions between two KASP markers might not have been detected.

### 4.2 Homozygous *T. timopheevii* introgressions in the wheat background

Homozygous introgressions from all linkage groups of *T. timopheevii* were obtained (Figures 3 and 4) covering 98.9% of the *T. timopheevii* genome. These introgressions covered the entire linkage groups of most At and G subgenomes except chromosomes 3A^t^, 4G, and 6G. However, more than 92% of these three chromosomes was introgressed into wheat. In total, 163 unique introgressions were observed, 89 (55%) from the A^t^ subgenome and 74 (45%) from the G subgenome. This was expected as previous reports have shown A^t^ and A genome chromosomes pair more readily than G and B genome chromosomes (Feldman 1966).

The highest number of unique homozygous introgressions were recovered from 5A^t^ (25) and 2G (20) which were present in 47 and 33 ILs, respectively, more than any other chromosome from the respective subgenomes. The high number of introgressions was expected for 2G as this chromosome shows a higher level of transmission through the gametes than expected (Devi et al., 2019; Nyquist 1962; Timonova et al., 2012). Preferential transmission of chromosome 2S of *Ae. speltoides*, proposed to be more closely related to the *T. timopheevii* G genome than to the B genome of wheat (Li et al., 2022; Rodriguez et al., 2000a), is well documented (King et al., 2018; Tsujimoto and Tsunewaki 1988) although the gene causing the preferential transmission in *T. timopheevii* appears to be weaker than the gene for preferential transmission in *Ae. speltoides*. A high incidence of introgressions from *T. timopheevii* linkage groups 1A^t^, 2A^t^, 2G, 5A^t^, 5G and 6G was also found in lines screened by Leonova et al., (2011) which when taken forward by Timonova et al., 2013 also showed a high transmission of the 2G segment. Analysis of the recent wheat pangenome also showed the presence of a large 2G introgression in chromosome 2B of wheat cultivar LongReach Lancer conferring stem rust resistance (Walkowiak et al., 2020).

The smallest number of introgressions was obtained for 4A^t^ (2 introgressions and a whole chromosome substitution). Recombination between chromosomes 4A of wheat and 4A^t^ of *T. timopheevii* is likely to have been reduced due to translocation differences between the two chromosomes. Although both carry the 4AL/5AL translocation, inherited from the A genome progenitor *T. urartu*, 4A of wheat has undergone additional paracentric and pericentric inversions (Devos et al., 1995; Dvorak et al., 2018) which are not present in *T. timopheevii* (Maestra and Naranjo 1999). These structural differences would result in reduced levels of recombination between the two chromosomes resulting in fewer introgressions.

A reduction in the number of introgressions was also expected from *T. timopheevii* chromosomal regions of 3A^t^, 6A^t^, 1G and 4G as four species-specific translocations involving these chromosomes occur within the *T. timopheevii* groups; 6A^t^S/1GS, 1GS/4GS, 4GS/4A^t^L and 4A^t^L/3A^t^L, plus a paracentric inversion in 6A^t^L (Jiang and Gill 1994; Maestra and Naranjo 1999; Rodriguez et al., 2000b; Salina et al., 2006). Chromosomes 3A and 3A^t^ have also been reported to have diverged through mutations in their primary DNA structure (Dobrovolskaya et al., 2009). From the regions involved in species-specific translocations two introgressions each were obtained for 3AL (3At.A9 and 3At.A10), 1GS (1G.B1 and 1G.B2) and 6AL (6At.A3 and 6At.A4) and one each for 6A^t^S (6At.A2) and 4GS (4G.B1; Figures 3 and 4).

A number of authors have also reported different accessions of *T. timopheevii* as carrying karyotype rearrangements (Badaeva et al., 1994; Badaeva et al., 1990; Badaeva et al., 2022; Kawahara and Tanaka 1977; Taketa and Kawahara 1996). Thus, in addition to the species-specific translocations described above, the accession used in this work may have carried additional rearrangements affecting recombination and hence introgression. This could also explain the differences in levels of recombination not just between chromosomes, e.g., the small number of introgressions from 7G, but between chromosome arms as seen in Figure 3. For example, recombination has occurred along the whole length of chromosomes 2A^t^/2A, 7A^t^/7A, 2B/2G and 3B/3D/3G (although lower in the pericentric regions), while recombination between 1A^t^/1A, 5A^t^/5A and 5G/5B has occurred more frequently in the long arm.

Due to high levels of synteny between A^t^ and A, and G and B genomes (Dvořák et al., 1993; Tsunewaki et al., 1996), the KASP markers validated for A and B genome chromosomes of wheat also, in majority of the cases, detected A^t^ and G chromosome segments, respectively (Table 1; Figure 2). The closeness of the two groups of genomes also resulted in the majority of A^t^ and G subgenome introgressions recombining with the A and B genomes of wheat, respectively. Although, 2% (2 out of 98) of the introgressions from the A^t^ subgenome and 26% (19 out of 74) of the introgressions obtained for the G subgenome of *T. timopheevii* had recombined with the D genome of wheat (Figures 3 and 4). It is possible that synteny between regions of the G subgenome and regions of the wheat D genome may be greater than between the G and B genomes especially for those chromosomes where either species-specific translocations or accession-specific translocations have occurred in *T. timopheevii*.

The At.D and G.D introgressions were validated through mcGISH (Figure 5A-D) since the D genome chromosomes can be easily distinguished against the A^t^ and G subgenomes of *T. timopheevii* unlike the A and B genomes of wheat. Due to the high sequence similarity between the A^t^ and A and G and B subgenomes, the mcGISH probes cannot distinguish between these groups of chromosomes. This meant that the At.A and G.B introgressions produced in this study could not be visualised using mcGISH. Instead, a few of these introgressions were validated using mcFISH (Figure 5E-H). Previous reports have done extensive karyotyping of all *T. timopheevii* chromosomes using mcFISH (Badaeva et al., 2016; Devi et al., 2019), however, characterising interspecific introgression using mcFISH is very laborious and time-consuming. This highlights the utility of the chromosome-specific KASP markers developed in this work which are capable of detecting the presence and site of *T. timopheevii* introgressions in wheat.

In addition to the 99 homozygous wheat-*T. timopheevii* ILs generated in this work (Figure 1; Table 2), further ILs have been generated within the Wheat Research Centre at Nottingham. With the availability of a high density of KASP markers, these ILs will be screened for unique introgressions from *T. timopheevii* with particular emphasis on those linkage groups from which relatively few introgressions have so far been obtained. The lines described in this paper are being screened for a variety of traits including Fusarium head blight resistance (unpublished results). Seed of lines Tim1-15, Tim17-19 and Tim21-35 have been deposited at the Germplasm Resource Unit at the John Innes Centre (https://www.seedstor.ac.uk/), while lines Tim36-99 will be distributed after seed multiplication.

## 6 Author Contributions

Germplasm generation, C.Y., S.A., D.S., S.H-E., S.G., I.P.K and J.K.; DNA extraction, C.Y., D.S. and S.A.; KASP marker development and genotyping, S.G, S.A., D.S., M.O. and J.W.; *in situ* hybridisation analysis, C.Y., M.O. and N.W.; Sequencing and SNP Discovery, B.C and A.H; Data analysis and curation, S.G.; Conceptualization, I.P.K., J.K and S.G.; manuscript writing, J.K., S.G, and I.P.K. and funding acquisition, I.P.K. and J.K. All authors have read and agreed to the published version of the manuscript.

## 8 Data availability

Raw reads data for *T. timopheevii* has been made available through the Grassroots data repository hosted by the Earlham Institute and funded by DFW programme (https://opendata.earlham.ac.uk/wheat). The genotyping data is available from the corresponding authors on reasonable request.

## 9 Supplemental Data

The supplemental tables provide detailed lists of all the chromosome-specific SNPs, between wheat and *T. timopheevii*, discovered in this work (Table S1), full set of validated chromosome-specific KASP markers that are detect subgenomes of *T. timopheevii* in a wheat background (Table S2) and KASP markers diagnostic for every introgression in each of the introgression lines (Table S3).

